## Supplementary Data for "Continuous millisecond conformational cycle of a DEAH box helicase reveals control of domain motions by atomic-scale transitions"

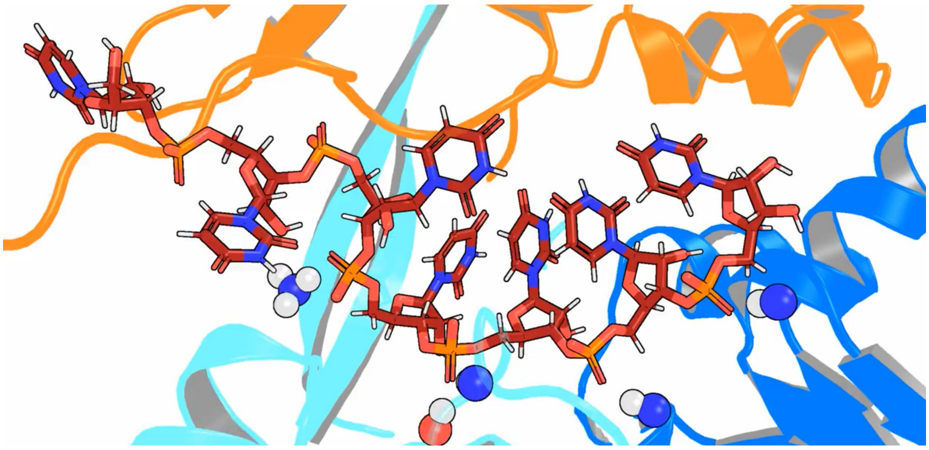

FIG. 1. Placeholder: Movie of the dynamics of RNA during the complete process.

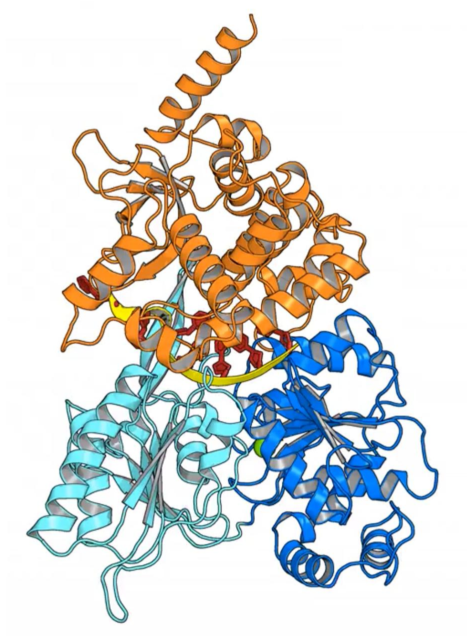

FIG. 2. Placeholder: Movie of the concatenated opening and closing process of the complete enzyme.

| | RecA-Dist. | G349-U5 | T381-U5 | R435-ATP | K403-U4 | E316-U4 | S387-Mg | S387-U5 | G349-U4-O1 | G349-U4-O2 | T381-U4-O1 | T381-U4-O2 | R180-H | R153-H | R180-C-U7 | K403-U3 | S387-Phi | S387-Psi |
|---|---|---|---|---|---|---|---|---|---|---|---|---|---|---|---|---|---|---|
| E-value | 2.65 | 0.7 | 0.81 | 0.38 | 1.0 | 0.7 | 0.26 | 1.58 | 0.17 | 0.32 | 0.16 | 0.4 | 0.18 | 0.19 | 0.9 | 0.33 | 59.0 | -47.0 |
| S-value | 3.1 | 0.19 | 0.26 | 0.75 | 0.29 | 0.21 | 1.44 | 0.2 | 0.92 | 0.73 | 0.9 | 0.73 | 0.83 | 0.62 | 0.48 | 0.87 | -70.0 | -40.0 |
| run1 | 2.7 | 0.63 | 0.75 | 0.4 | 1.01 | 0.82 | 0.52 | 1.23 | 0.18 | 0.33 | 0.17 | 0.41 | 0.23 | 0.18 | 1.04 | 0.27 | -131.14 | 149.37 |
| run2 | 2.74 | 0.64 | 0.81 | 0.39 | 0.9 | 0.8 | 0.5 | 1.21 | 0.19 | 0.31 | 0.18 | 0.42 | 0.2 | 0.19 | 1.05 | 0.28 | -161.8 | 154.18 |
| run3 | 2.71 | 0.6 | 0.8 | 0.38 | 0.96 | 0.78 | 0.56 | 1.11 | 0.18 | 0.33 | 0.18 | 0.41 | 0.2 | 0.19 | 1.06 | 0.27 | -151.21 | 150.41 |
| run4 | 2.73 | 0.62 | 0.78 | 0.39 | 0.89 | 0.82 | 0.49 | 1.22 | 0.18 | 0.31 | 0.18 | 0.43 | 0.2 | 0.18 | 1.05 | 0.27 | -160.6 | 153.45 |
| run5 | 2.75 | 0.62 | 0.82 | 0.4 | 1.03 | 0.78 | 0.5 | 1.16 | 0.18 | 0.32 | 0.17 | 0.42 | 0.2 | 0.19 | 1.05 | 0.28 | -148.62 | 152.54 |
| run6 | 2.77 | 0.63 | 0.75 | 0.4 | 0.95 | 0.83 | 0.56 | 1.23 | 0.19 | 0.31 | 0.18 | 0.41 | 0.21 | 0.18 | 1.03 | 0.27 | -162.34 | 147.8 |
| run7 | 2.76 | 0.61 | 0.77 | 0.39 | 0.96 | 0.8 | 0.52 | 1.14 | 0.18 | 0.32 | 0.17 | 0.42 | 0.2 | 0.19 | 1.06 | 0.27 | -148.18 | 150.75 |
| run8 | 2.72 | 0.63 | 0.79 | 0.39 | 0.9 | 0.82 | 0.51 | 1.23 | 0.19 | 0.31 | 0.18 | 0.42 | 0.22 | 0.19 | 1.04 | 0.28 | -161.9 | 150.54 |
| run9 | 2.74 | 0.62 | 0.78 | 0.4 | 0.94 | 0.8 | 0.5 | 1.23 | 0.18 | 0.32 | 0.18 | 0.42 | 0.21 | 0.18 | 1.02 | 0.27 | -147.2 | 152.26 |
| run10 | 2.73 | 0.63 | 0.8 | 0.4 | 0.9 | 0.83 | 0.52 | 1.25 | 0.19 | 0.3 | 0.18 | 0.43 | 0.21 | 0.18 | 1.0 | 0.28 | -163.25 | 150.81 |
| run11 | 2.74 | 0.6 | 0.8 | 0.39 | 0.93 | 0.8 | 0.53 | 1.12 | 0.18 | 0.31 | 0.18 | 0.42 | 0.2 | 0.19 | 1.03 | 0.27 | -164.92 | 150.29 |
| run12 | 2.73 | 0.6 | 0.79 | 0.38 | 0.96 | 0.83 | 0.54 | 1.13 | 0.18 | 0.32 | 0.18 | 0.42 | 0.21 | 0.18 | 1.06 | 0.28 | -130.48 | 138.06 |
| run13 | 2.72 | 0.64 | 0.78 | 0.4 | 0.94 | 0.77 | 0.51 | 1.23 | 0.18 | 0.33 | 0.18 | 0.42 | 0.22 | 0.18 | 1.05 | 0.28 | -160.61 | 148.87 |
| run14 | 2.76 | 0.64 | 0.8 | 0.39 | 0.92 | 0.81 | 0.56 | 1.25 | 0.19 | 0.32 | 0.17 | 0.42 | 0.2 | 0.18 | 0.99 | 0.27 | -145.06 | 151.91 |
| run15 | 2.72 | 0.61 | 0.78 | 0.4 | 0.94 | 0.81 | 0.53 | 1.24 | 0.18 | 0.32 | 0.17 | 0.41 | 0.2 | 0.18 | 1.06 | 0.28 | -112.83 | 149.76 |
| run16 | 2.72 | 0.63 | 0.84 | 0.38 | 0.95 | 0.81 | 0.55 | 1.12 | 0.19 | 0.3 | 0.17 | 0.39 | 0.2 | 0.19 | 1.05 | 0.28 | -162.92 | 157.06 |
| run17 | 2.72 | 0.61 | 0.8 | 0.39 | 0.94 | 0.8 | 0.54 | 1.09 | 0.18 | 0.32 | 0.18 | 0.42 | 0.21 | 0.18 | 1.05 | 0.27 | -166.71 | 151.9 |
| run18 | 2.74 | 0.61 | 0.79 | 0.4 | 0.89 | 0.8 | 0.49 | 1.24 | 0.18 | 0.32 | 0.18 | 0.43 | 0.2 | 0.19 | 1.07 | 0.27 | -148.49 | 147.3 |
| run19 | 2.73 | 0.62 | 0.82 | 0.4 | 0.93 | 0.77 | 0.54 | 1.09 | 0.18 | 0.33 | 0.18 | 0.42 | 0.21 | 0.2 | 1.03 | 0.27 | -131.54 | 149.69 |
| run20 | 2.74 | 0.63 | 0.78 | 0.4 | 0.95 | 0.76 | 0.53 | 1.08 | 0.18 | 0.3 | 0.18 | 0.42 | 0.21 | 0.19 | 1.04 | 0.28 | -150.95 | 150.79 |
| run21 | 2.72 | 0.64 | 0.82 | 0.39 | 0.93 | 0.8 | 0.52 | 1.09 | 0.19 | 0.31 | 0.17 | 0.41 | 0.2 | 0.19 | 1.05 | 0.28 | -163.88 | 151.64 |
| run22 | 2.73 | 0.61 | 0.78 | 0.39 | 0.92 | 0.81 | 0.53 | 1.08 | 0.18 | 0.32 | 0.17 | 0.42 | 0.21 | 0.18 | 1.07 | 0.28 | -164.43 | 144.9 |
| run23 | 2.73 | 0.62 | 0.79 | 0.39 | 0.83 | 0.76 | 0.52 | 1.13 | 0.19 | 0.31 | 0.18 | 0.43 | 0.21 | 0.18 | 1.05 | 0.28 | -164.11 | 151.2 |
| run24 | 2.74 | 0.74 | 0.88 | 0.4 | 0.92 | 0.81 | 0.49 | 1.2 | 0.35 | 0.23 | 0.18 | 0.29 | 0.21 | 0.18 | 1.07 | 0.28 | -163.43 | 149.93 |
| run25 | 2.72 | 0.63 | 0.79 | 0.38 | 0.88 | 0.74 | 0.56 | 1.1 | 0.19 | 0.34 | 0.18 | 0.41 | 0.2 | 0.19 | 1.04 | 0.28 | -116.97 | 146.64 |
| run26 | 2.72 | 0.61 | 0.79 | 0.4 | 0.89 | 0.83 | 0.52 | 1.12 | 0.19 | 0.3 | 0.18 | 0.42 | 0.21 | 0.18 | 1.06 | 0.28 | -166.16 | 140.06 |
| run27 | 2.71 | 0.69 | 0.78 | 0.4 | 0.82 | 0.8 | 0.53 | 1.24 | 0.19 | 0.32 | 0.18 | 0.43 | 0.21 | 0.18 | 1.05 | 0.28 | -162.65 | 150.29 |
| run28 | 2.74 | 0.61 | 0.77 | 0.4 | 0.92 | 0.78 | 0.53 | 1.23 | 0.18 | 0.32 | 0.18 | 0.43 | 0.21 | 0.18 | 1.05 | 0.28 | -148.24 | 150.09 |
| run29 | 2.71 | 0.73 | 0.88 | 0.38 | 0.92 | 0.84 | 0.53 | 1.03 | 0.32 | 0.24 | 0.17 | 0.31 | 0.2 | 0.18 | 1.05 | 0.28 | -131.87 | 146.6 |
| run30 | 2.74 | 0.65 | 0.81 | 0.4 | 0.88 | 0.78 | 0.56 | 1.24 | 0.19 | 0.3 | 0.18 | 0.43 | 0.21 | 0.18 | 1.04 | 0.28 | -149.43 | 150.91 |
| run31 | 2.71 | 0.64 | 0.77 | 0.4 | 0.96 | 0.82 | 0.53 | 1.25 | 0.18 | 0.31 | 0.17 | 0.42 | 0.21 | 0.18 | 0.99 | 0.28 | -158.86 | 150.5 |
| run32 | 2.72 | 0.6 | 0.78 | 0.38 | 0.89 | 0.82 | 0.53 | 1.19 | 0.18 | 0.31 | 0.18 | 0.43 | 0.2 | 0.19 | 0.99 | 0.28 | -149.01 | 152.9 |
| run33 | 2.73 | 0.59 | 0.79 | 0.4 | 0.91 | 0.76 | 0.52 | 1.11 | 0.18 | 0.32 | 0.17 | 0.42 | 0.21 | 0.19 | 1.0 | 0.27 | -148.76 | 149.11 |
| run34 | 2.77 | 0.64 | 0.78 | 0.4 | 0.89 | 0.83 | 0.54 | 1.24 | 0.19 | 0.33 | 0.19 | 0.43 | 0.21 | 0.18 | 1.03 | 0.27 | -149.98 | 145.62 |
| run35 | 2.73 | 0.62 | 0.81 | 0.39 | 0.93 | 0.79 | 0.55 | 1.07 | 0.18 | 0.33 | 0.17 | 0.42 | 0.21 | 0.19 | 1.03 | 0.28 | -164.43 | 141.47 |
| run36 | 2.75 | 0.64 | 0.82 | 0.39 | 0.95 | 0.76 | 0.52 | 1.08 | 0.18 | 0.32 | 0.17 | 0.41 | 0.2 | 0.19 | 1.06 | 0.27 | -165.8 | 148.06 |
| run37 | 2.73 | 0.64 | 0.83 | 0.38 | 0.96 | 0.76 | 0.55 | 1.11 | 0.22 | 0.3 | 0.17 | 0.38 | 0.21 | 0.2 | 1.07 | 0.28 | -144.23 | 130.64 |
| run38 | 2.71 | 0.62 | 0.77 | 0.39 | 0.99 | 0.78 | 0.51 | 1.25 | 0.18 | 0.32 | 0.17 | 0.42 | 0.21 | 0.19 | 1.03 | 0.27 | -145.39 | 151.66 |
| run39 | 2.67 | 0.62 | 0.81 | 0.38 | 0.92 | 0.75 | 0.55 | 1.09 | 0.18 | 0.32 | 0.17 | 0.41 | 0.2 | 0.19 | 1.04 | 0.27 | -164.67 | 148.58 |
| run40 | 2.75 | 0.71 | 0.87 | 0.39 | 0.9 | 0.79 | 0.53 | 1.16 | 0.33 | 0.24 | 0.17 | 0.3 | 0.21 | 0.18 | 1.07 | 0.28 | 36.3 | 61.5 |
| run41 | 2.76 | 0.65 | 0.8 | 0.38 | 0.89 | 0.81 | 0.53 | 1.06 | 0.18 | 0.32 | 0.18 | 0.43 | 0.22 | 0.19 | 1.04 | 0.27 | -119.08 | 143.61 |
| run42 | 2.74 | 0.67 | 0.82 | 0.38 | 0.81 | 0.76 | 0.53 | 1.13 | 0.22 | 0.27 | 0.17 | 0.41 | 0.21 | 0.19 | 1.03 | 0.28 | -98.64 | 147.45 |
| run43 | 2.74 | 0.6 | 0.79 | 0.39 | 0.95 | 0.8 | 0.53 | 1.18 | 0.18 | 0.32 | 0.17 | 0.42 | 0.21 | 0.19 | 1.05 | 0.27 | -146.06 | 146.37 |
| run44 | 2.74 | 0.62 | 0.79 | 0.4 | 0.91 | 0.84 | 0.49 | 1.24 | 0.18 | 0.29 | 0.18 | 0.43 | 0.2 | 0.18 | 1.02 | 0.27 | -131.91 | 145.61 |
| run45 | 2.67 | 0.62 | 0.79 | 0.39 | 0.91 | 0.84 | 0.53 | 1.19 | 0.18 | 0.31 | 0.17 | 0.42 | 0.2 | 0.18 | 0.99 | 0.28 | -162.12 | 149.31 |
| run46 | 2.73 | 0.61 | 0.78 | 0.38 | 0.94 | 0.79 | 0.53 | 1.08 | 0.18 | 0.32 | 0.17 | 0.42 | 0.21 | 0.18 | 1.05 | 0.27 | -31.91 | 144.47 |
| run47 | 2.73 | 0.62 | 0.81 | 0.4 | 0.96 | 0.7 | 0.56 | 1.18 | 0.19 | 0.33 | 0.17 | 0.41 | 0.19 | 0.19 | 1.04 | 0.29 | -165.88 | 147.11 |
| run48 | 2.75 | 0.63 | 0.81 | 0.39 | 0.96 | 0.77 | 0.51 | 1.21 | 0.2 | 0.31 | 0.17 | 0.4 | 0.2 | 0.18 | 1.01 | 0.28 | -165.3 | 150.54 |
| run49 | 2.73 | 0.65 | 0.83 | 0.39 | 0.93 | 0.8 | 0.54 | 1.12 | 0.18 | 0.33 | 0.18 | 0.42 | 0.22 | 0.2 | 1.06 | 0.27 | -147.62 | 151.2 |
| run50 | 2.73 | 0.64 | 0.81 | 0.4 | 0.88 | 0.83 | 0.51 | 1.23 | 0.19 | 0.3 | 0.2 | 0.45 | 0.23 | 0.18 | 1.08 | 0.28 | -114.69 | 150.83 |

FIG. 3. Example heat table from the closing process, visualizing the progression of one round of AS simulations towards the target structure.

Rows correspond to 50 individual simulations of one round of AS. Top row: list of structural features including distances, angles, and $\phi/\psi$ angles. Second row: reference (target) values of the features, here taken from the 6I3P structure of Prp43•$U_7$•ATP. Third row: starting values of the features, taken from the open simulation frame. Columns show the feature values at the end of this AS round. The color indicates the similarity with the starting feature (purple) or with the reference/target feature (yellow). Such tables have been used extensively to monitor the progression of the AS simulations and to select the most successful simulation to be used a seed for the next round of AS.

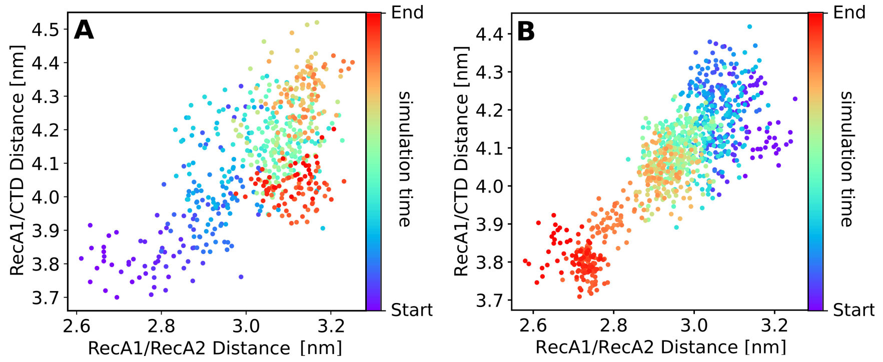


FIG. 4. Center-of-mass distances between C-terminal domain relative to RecA-like domains. (A) RecA1–CTD distance versus RecA1–RecA2 distance during opening and (B) during closing. Here, the center of mass of CTD was defined with the helices S544–S555, D583–R600, and Y615–K629. The color indicates the cumulative simulation time from start (purple) to end (red). During both the opening (A) and (B) closing process, the RecA1–CTD distance was correlated with the RecA1–RecA2 distance, illustrating that the CTD domain moved concertedly with RecA2 along the RNA, as shown in fig. 2 in the paper.

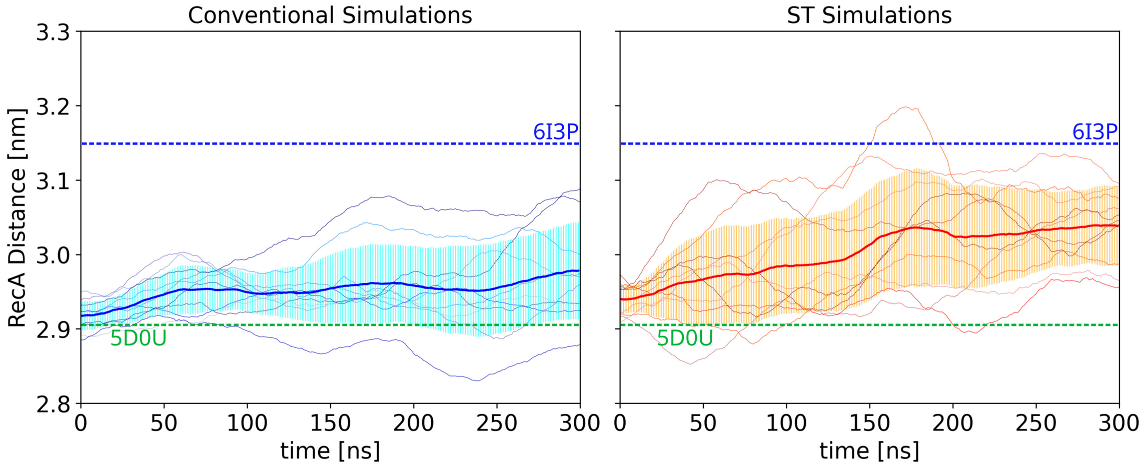


FIG. 5. Accelerated conformational sampling with simulated tempering (ST). RecA1–RecA2 distance of Prp43 after removal of ADP in conventional simulations (left) or ST simulations (right). Among 10 conventional simulations, only two simulation reached a semi-open state within 300 ns, as indicated by the RecA1/RecA2 distance of over 3.0 nm. In contrast, among 10 ST simulations, four reached the semi-open state within only 100 ns and six simulation reached the semi-open state within 300 ns. In addition, four ST reached a fully open state indicated by a distance larger 3.1 nm. The more rapid increase of the average (thick lines) and of standard deviations (shaded areas) in ST compared to conventional simulations indicate accelerated sampling of the conformational space within a short simulation time.

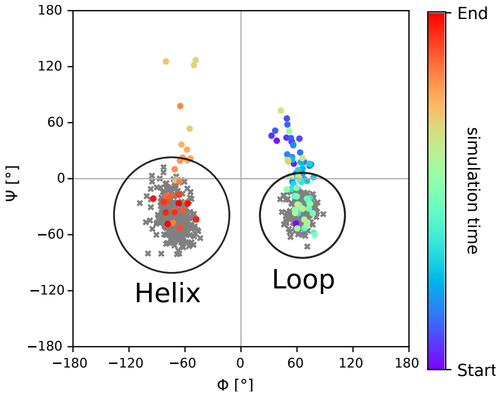


FIG. 6. Ramachandran plot of sensor serine S387.

Grey crosses: $\phi$ and $\psi$ angles of S387 during free 1 $\mu$s simulations starting from the 6I3P structure (left), corresponding to the open conformation, or starting from the 5LTA structure (right), corresponding to the closed conformation. According to the $\phi/\psi$ angles, S387 remained in the helical or in a loop state in the simulations of the open or closed state, respectively. No transition was observed. Colored dots: $\phi/\psi$ angles during a AS simulation with a successful loop-to-helix transition of the sensor serine. The color indicates the simulation time of the successful AS run from purple to red.

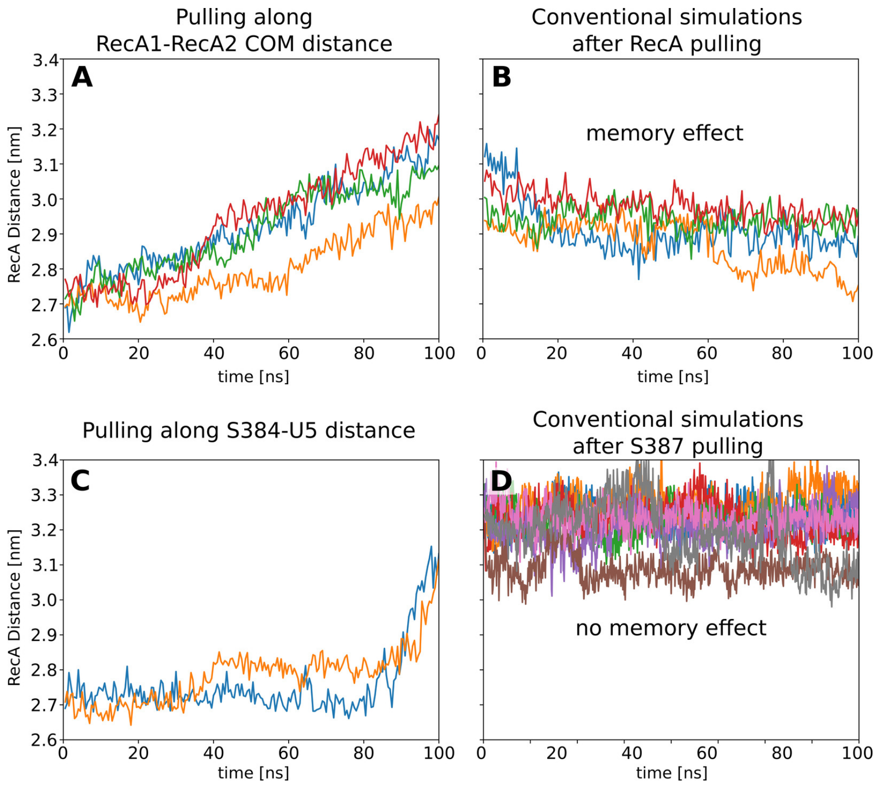


FIG. 7. On the correlation between Prp43 opening and loop-to-helix transition of the S387–G392 sensor loop.

RecA1–RecA2 distance versus simulation time during (A/C) pulling simulations and (B/D) after releasing the pulling restraint. (A) Pulling along the RecA1–RecA2 center-of-mass (COM) distance leads to major memory effects, as shown by (B) the partial re-closure of the RecA1/RecA2 interface after release of the restraint. (C) Pulling along the S387–U5 distance, thereby driving the loop-to-helix transition of the sensor loop, leads to opening of the RecA1–RecA2 interface and (D) does not lead to memory effects. The absence of memory effects after the loop-to-helix transition suggests that the sensor loop transitions are critical for Prp43 opening and closing.

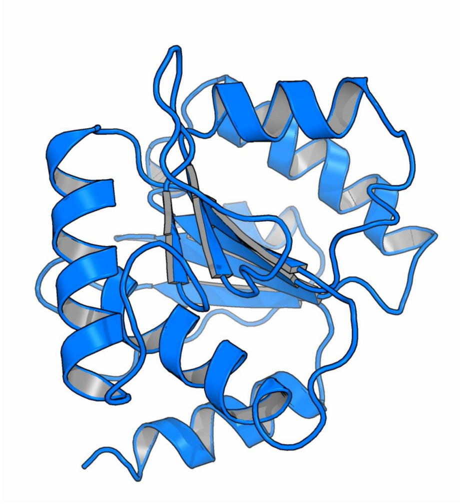

FIG. 8. Placeholder: Movie of the projected first eigenvector of the PCA of the isolated RecA1 domain.

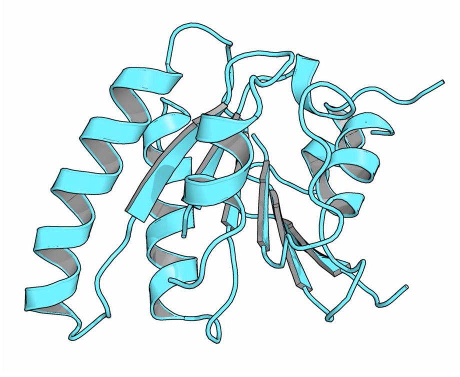

FIG. 9. Placeholder: Movie of the projected first eigenvector of the PCA of the isolated RecA2 domain.

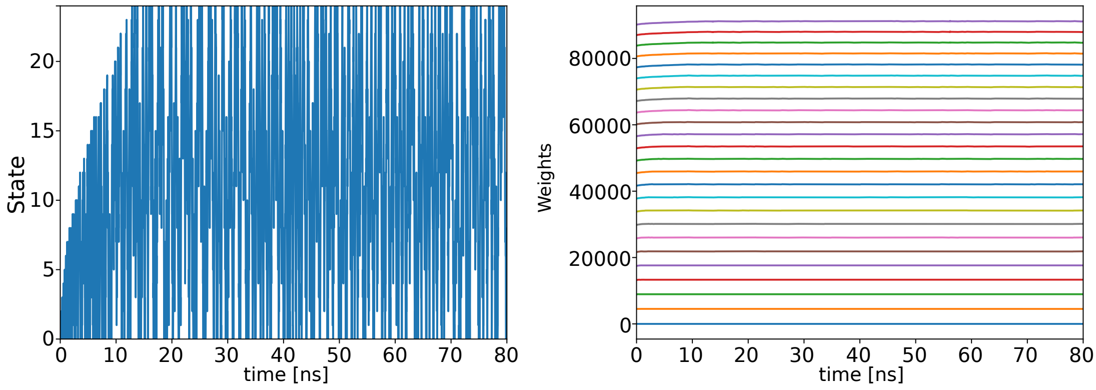


FIG. 10. Coverage of temperature states during simulated tempering.
Left: temperature state versus simulation time during an example simulated tempering (ST) simulation. During the burn-in phase within the first 17 ns, states with increasingly higher temperature were visited, reflecting the gradual adaptation of the weights. Right: weights of temperature states versus simulation time. Weights were only slightly adapted at the beginning of the simulation, reflecting good initial weights. After 17 ns, the weights were converged, and all states were frequently visited.

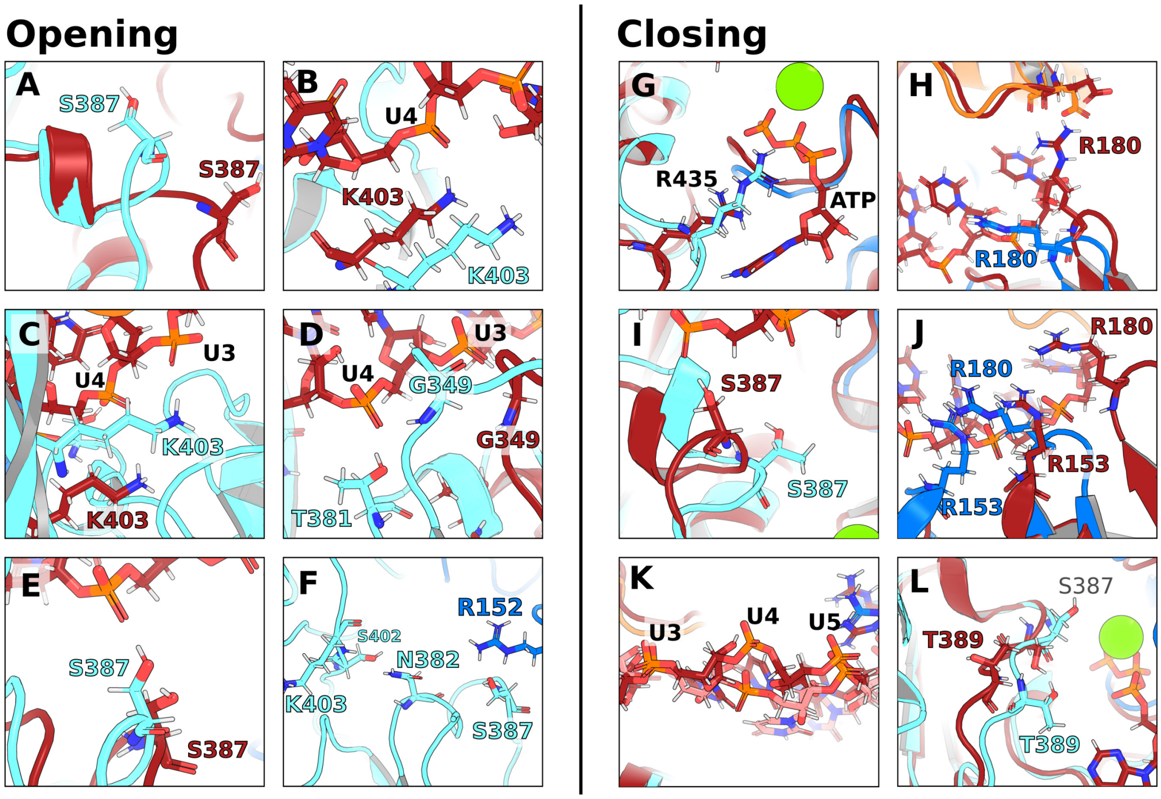


FIG. 11. Transitions of molecular switches contributing to the mean first passage times. Opening: (A) 1st transition: S387 flip from loop (red) to helical state (cyan). (B) 2nd transition: K403 connected to U4 (red) to disconnected to U4 (cyan). (C) 3rd transition: K403 disconnected to U3 (red) to connected to U3 (cyan). (D) 4th transition: Hook-loop shift from U3 (G349; red) to U4 (G349 and T381; cyan). (E) 5th transition: S387 not connected to U5 (red) to S387 connected to U5 (cyan). (F) 6th transition: H-bond network after successful opening. RecA2 is connected with RecA1 with an H-bond between R152 and S387. The N382 of the sensor loop is connected with the $\beta$-hairpin via K403 and S402. Closing: (G) 1st transition: R435 binding to $\beta$ and $\gamma$ phosphates of ATP (from red to cyan). (H) 2nd transition: R180 bound to CTD (red) to disconnected from CTD (blue). (I) 3rd transition: S387 flip from helical (red) to loop (cyan) state. (J) 4th transition: Hook-turn shift from U6/U7 (red) to U5/U6 (blue). (K) 5th transition: RNA rotation at U4 (from pink to red). (L) 6th transition: Bending of S387 and T389 during the end of the closing process (from red to cyan), finalizing the formation of the RecA1/RecA2 interface corresponding to a RecA distance of 2.65 nm.

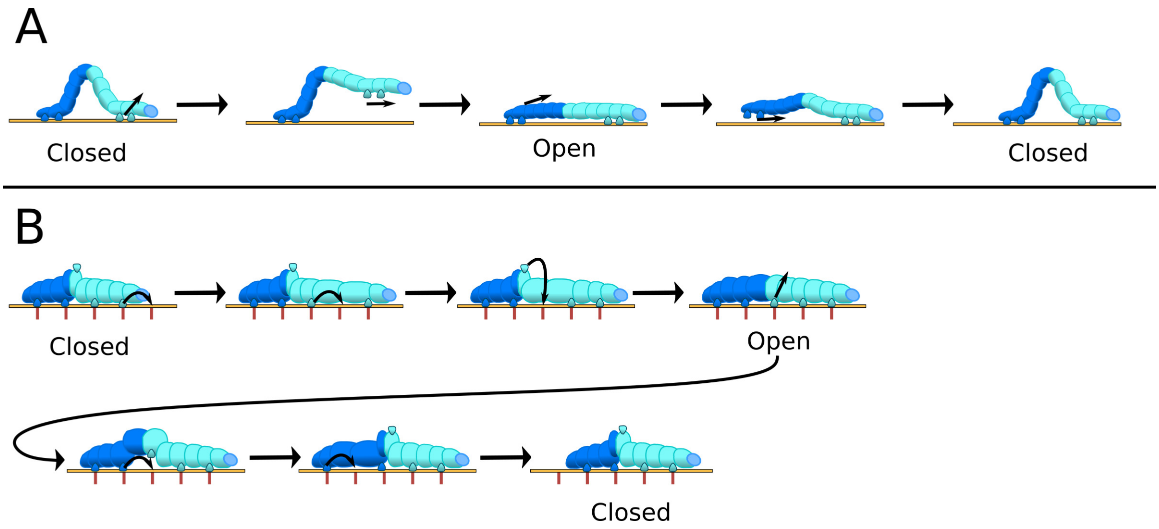


FIG. 12. Classical inchworm model and proposed inchworm/caterpillar model.

A: The classical inchworm model may imply that RecA-like domains fully detach from the RNA strand one after another, similar to the leg movement of an inchworm. B: Proposed inchworm/caterpillar model to illustrate both the center of mass motion of the RecA-like domains (dark blue and cyan) and the crawling of RecA-like along the RNA. Hydrogen bond partners of protein and RNA are sketched as caterpillar legs and as red lines, respectively. Sine the RecA-like domains bind the RNA with four or five H-bonds in the closed and open state, respectively, the caterpillar requires five legs. The central leg, modeling the sensor serine S387, carries out a rotation to bind the free nucleotide binding site before reaching the open state.

TABLE I. Key properties of the adaptive sampling rounds.

For each round of adaptive sampling, sum of simulation times $t_{\mathrm{sim}}$, number of simulations $N_{\mathrm{sim}}$, number of observed transitions $N_{\mathrm{trans}}$, estimated transition rate $k$ for the opening (left) and closing process (right).

| | Opening | | | | Closing | | | |
|---|---|---|---|---|---|---|---|---|
| AS round | $t_{\mathrm{sim}}$ (µs) | $N_{\mathrm{sim}}$ | $N_{\mathrm{trans}}$ | $k$ (1/µs) | $t_{\mathrm{sim}}$ (µs) | $\mathrm{N}_{\mathrm{sim}}$ | $N_{\mathrm{trans}}$ | $k$ (1/µs) |
| 1 | 10.0 | 100 | 1 | 0.10 | 0.8 | 10 | 3 | 3.69 |
| 2 | 10.0 | 100 | 1 | 0.10 | 0.7 | 10 | 0 | – |
| 3 | 10.0 | 100 | 9 | 0.90 | 0.7 | 10 | 0 | – |
| 4 | 10.0 | 100 | 0 | – | 2.0 | 30 | 1 | 0.29 |
| 5 | 5.0 | 500 | 2 | 0.13 | 3.0 | 30 | 0 | – |
| 6 | 8.3 | 100 | 20 | 2.41 | 2.0 | 30 | 2 | 0.40 |
| 7 | 0.4 | 40 | 0 | – | 2.0 | 20 | 2 | 1.00 |
| 8 | 0.3 | 40 | 0 | – | 4.0 | 40 | 0 | – |
| 9 | 1.6 | 40 | 1 | 0.43 | 5.0 | 50 | 5 | 0.56 |
| 10 | – | – | – | – | 5.0 | 50 | 0 | – |
| 11 | – | – | – | – | 4.0 | 40 | 0 | – |
| 12 | – | – | – | – | 3.5 | 40 | 0 | – |
| 13 | – | – | – | – | 2.7 | 40 | 1 | 0.06 |
